# Compensation and conversation in autism: atypical connectivity supports typical behavior

**DOI:** 10.1101/2022.11.18.517079

**Authors:** Kyle Jasmin, Alex Martin, Stephen J. Gotts

## Abstract

**Background:** It is well-established that individuals with autism show atypical functional brain connectivity. However, the role this plays in behavior, especially in naturalistic social settings, has remained unclear. Some atypical patterns may reflect core deficits, while others may instead compensate for deficits and promote adaptive behavior. Distinguishing these possibilities requires measuring the ‘typicality’ of spontaneous behavior and determining which connectivity patterns correlate with it.

**Methods:** Thirty-nine male participants (19 autism, 20 typically-developed) engaged in 115 spontaneous conversations with an experimenter during fMRI scanning (Jasmin, et al., 2019, Brain). A classifier algorithm was trained to distinguish participants by diagnosis based on 81 semantic, affective and linguistic dimensions derived from their use of language. The algorithm’s certainty that a participant was in either the autism or typical group was used as a measure of task performance and compared with functional connectivity levels.

**Results:** The algorithm accurately classified participants (74%, *P* = .002), and its scores correlated with clinician-observed autism signs (ADOS) (*r_s_* = .56, *P* = .03). In support of a compensatory role, greater functional connectivity, most prominently between left-hemisphere social communication regions and right inferior frontal cortex, correlated with more typical language behaviour, only for the autism group (*r*_s_ = .56, *P* = .01).

**Conclusion:** We report a simple and highly generalizable method for quantifying behavioral performance and neural compensation during complex spontaneous social behavior, without the need for an *a priori* benchmark. The findings suggest that functional connectivity increases in autism during communication reflect a neural compensation strategy.

## Introduction

Neural studies of autism have often used functional connectivity analyses to identify networks that are under- or over-connected relative to neurotypical controls. Many of these studies have assessed connectivity during the resting state, and have identified both decreases (especially among cortical social processing areas), and increases (especially involving subcortical structures) (Di Martino et al., 2014; Gotts et al., 2012). Other studies have examined connectivity during social or language tasks, which also report both decreases (Just et al., 2004; Kana et al., 2006) and increases (Shen et al., 2012) in connectivity. However, the relationship between atypical connectivity and behavior has remained unclear. Some atypical patterns may cause social impairments (see (Gotts et al., 2019)). Others, especially those that emerge only during tasks, might instead reflect alternative neural strategies that compensate for core deficits (Johnson et al., 2015; Livingston & Happé, 2017). Determining which patterns promote adaptive behavior, and which ones hinder it will be crucial for understanding autism more fully and developing connectivity-based interventions (Gotts et al., 2019; Ramot et al., 2017).

One common technique for assessing the role of a given network in behavior has been to test for correlations with some measure of symptom severity, such as the Social Responsiveness Scale (Constantino et al., 2003), a questionnaire undertaken by parents or teacher that measures aspects of social cognition and autistic mannerisms. This tool has strong psychometric properties and may therefore accurately measure core autism-related deficits, as it is administered by someone (often a parent or teacher) who knows the individual well. However, tools like SRS that measure autistic traits may be insufficient for assessing behavioral compensation. This is because compensation reflects not the severity of core social deficits, but the degree to which overt behavior superficially resembles typical behavior in the face of these core deficits (Livingston & Happé, 2017). Assessing neural compensation in complex spontaneous social behavior therefore requires quantifying how typical behavior is during social interactions and then relating this measure to functional connectivity obtained during the same interactions. Evidence for compensation in autistic participants would consist of increasingly dissimilar functional connectivity relative to neurotypical controls in a relevant brain network as behavior becomes increasingly control-like.

We recently reported a study investigating face-to-face conversation in autism which makes it possible to examine compensatory behavior. Autism and typical participants engaged in spontaneous ‘face-to-face’ social interactions with an experimenter through video and audio links while being scanned with functional MRI (Jasmin et al., 2019). Measurements of neural activity during the task were compared with resting state. Some task and resting state results were similar, namely increased connectivity between subcortical structures (thalamus and ventral striatum) and the cortex. By contrast, the cortico-cortical pattern differed markedly between the resting and task states: during social interaction, widespread increases in connectivity, rather than decreases, were detected within a distributed, bilateral network of brain regions (Jasmin et al., 2019). We hypothesized that the overconnectivity may reflect a compensatory strategy to meet the demands of the difficult social task, but we lacked an objective measure of task performance that would have allowed us to test this account.

Here we now report an objective measure of the autism phenotype as it relates to language produced during the conversations in that experiment, which was developed by training a classifier algorithm on the transcribed speech of the autism and typical participants. The metric was validated by assessing how accurately it classified participants by diagnostic category and how well it correlated with clinician ratings on the ADOS-2, the ‘gold standard’ autism diagnostic interview tool (Lord et al., 2000). Finally, functional connectivity was assessed. If connectivity increases during conversation are compensatory, then elevated connectivity should correspond to a less pronounced autism phenotype during the interactions.

## Methods and Materials

### Participants

Nineteen males (aged 14.7 to 28.2 years) with autism and 20 male typically-developed participants (aged 15.1 to 32.0 years) took part. Participants with autism were recruited from the Washington, DC metropolitan area and met DSM-5 criteria for Autism Spectrum Disorder (APA, 2013) as assessed by an experienced clinician. All participants with autism received the ADOS-2 Module 4.(Lord et al., 2000) The scores from participants with autism met cut-off for the ‘broad autism spectrum disorders’ category according to criteria established by the National Institute of Child Health and Human Development/National Institute on Deafness and Other Communication Disorders Collaborative Programs for Excellence in Autism (Lainhart et al., 2006) The distributions for full-scale IQ, verbal IQ, and age did not differ statistically between the autism and typical groups (Jasmin et al., 2019). The experiment was approved by the NIMH Institutional Review Board (protocol 10-M-0027, clinical trials number NCT01031407)

### Procedure

Each session consisted of three spontaneous conversations between the participant and the experimenter. Prior to scanning, participants were told that they would engage in three unstructured and informal conversations. Using a questionnaire (Anthony et al., 2013), participants rated their level of interest in various topics such as music, games, and transportation vehicles, and indicated their top three interests, from which the experimenter selected two to serve as conversation topics. The topic of the final conversation was always work or school life, depending on participant’s age. The topics of conversations are listed in Supplementary Table 2 of Jasmin et al., 2019 (Jasmin et al., 2019). Before each conversation run, the experimenter sat in front of a blue screen facing a camera. The run began with 16 seconds of rest. Then, live video and audio from the experimenter were presented to the subject and the interaction task began. Conversations proceeded for 6 minutes. After each interaction, the video faded to black and a ‘STOP’ slide was displayed to the participant, followed by 30 additional seconds of rest to allow for delayed hemodynamic effects.

### Behavioral data processing

The audio recordings of the conversations were transcribed by professional transcriptionists and the text was analyzed with the Linguistic Inquiry Word Count (LIWC; pronounced “Luke”), 2007 Edition (Tausczik & Pennebaker, 2010). LIWC outputs 81 variables that reflect linguistic aspects of a text (e.g. total word count, number of words per sentence), as well as counts of words in particular linguistic, semantic and affective categories (e.g. affective words, sensory words, pronouns, articles).

The values of the LIWC output variables were z-scored, and a cross-validated linear support vector machine (SVM) model was trained on the 115 conversations using a leave-one-out approach (*CVSVMModel* function in MATLAB 2020b). The output scores of the left-out conversations were transformed using the *fitSVMPosterior* function to probabilities that reflected the certainty of the classification decision for each conversation (Platt, 1999). Lower scores indicated more certainty of autism classification according to the model, and higher scores indicated more certainty of typical classification. The probability scores for each conversation were then averaged together to yield a single composite score for each participant, which if lower than 0.5 indicated an overall classification of autism by the algorithm.

Performance of the classifier was assessed by computing accuracy (percent correct classifications relative to actual diagnosis). Statistical significance of classifier accuracy was determined by permutation test: a null distribution was compiled by permuting the diagnosis label for the participants, running a leave-one-out crossvalidation, and averaging the resulting probability scores by participant. The classifications derived from permuted data were compared to the actual clinical diagnostic status over 1000 permutations, and the reported *p*-value reflects the proportion of these permutations that resulted in accuracy that was better for the permuted labels than the actual labels.

External validity of the machine classifier measure was assessed by correlating the machine-derived classifier scores with clinician-derived ADOS-2 (Social + Communication) scores from the same participants. As a final control procedure, to exclude the possibility that any patterns of results were driven by systematic differences in experimenter behavior, another SVM classifier was trained on language produced by the experimenter during the same conversations with identical analysis pipeline and validation procedure applied. For an analysis of which LIWC categories most strongly drove the classification, see Supplementary Materials. As previously reported, gross measures of linguistic behavior of the autism and typical groups were similar: there were no statistical differences in the proportion of time that participants (vs. the experimenter) spent speaking [t(37) = 0.31, *p* = 0.76], the total number of words produced [t(37) = –1.4, *p* = 0.17] the count of speaking turns [t(37) = 0.46, *p* = 0.65], or the number of words per sentence [t(37) = 0.01, *p* = 0.99].

### MRI imaging and ROI selection

Having established a behavioral measure of typical language use, we then used a data-driven procedure to localize brain areas that showed the greatest functional connectivity in autism relative to the neurotypical control group. First, for each functional MRI conversation scan, a global connectivity map (or ‘whole-brain connectedness map’) was calculated. To do this, 1) each gray matter voxel was correlated with every other gray matter voxel, 2) those correlations were averaged together, and 3) the average correlation was stored back in the original voxel’s location (Cole et al., 2010; Gotts et al., 2012; Jasmin et al., 2020, 2019). Next, a contrast of Autism > Typical participants was performed on the connectedness maps after fitting a linear mixed effects model on the data with AFNI’s *3dLME*, controlling for Motion and Age (model = [Connectedness ~ Group + Age + Motion + (1 | Participant)].

Different combinations of voxel-wise significance thresholds and cluster extent thresholds result in different patterns of significant results. We sought to identify the brain areas most robust to choice of threshold by assessing voxel-wise and clusterwise significance across a range of thresholds. The Autism > Typical contrast was thresholded at P < .005, .001, .0005, .0001, .00005, and .00001, with corresponding cluster extent thresholds of *k* ≥ 51, 22, 15, 7, 5, and 2 (determined using AFNI’s *3dClustSim* with empirically derived spatial autocorrelation function, -acf; see Cox et al., 2017; Eklund et al., 2016)). The significance maps from each thresholding procedure were binarized (1 if significant, 0 otherwise) and combined to produce a map that illustrated the maximum threshold survived by each voxel). Only three regions showed significantly greater whole-brain functional connectivity in autism at every threshold tested: right mid-superior temporal sulcus (mSTS), right anterior STS, and right IFG/operculum (Figure 1; Table S1). These regions were selected for further analysis (at *P* < .0001, corrected, where each had 10 or more voxels, mitigating effects of between-subject anatomical variability).

**Figure 1.**
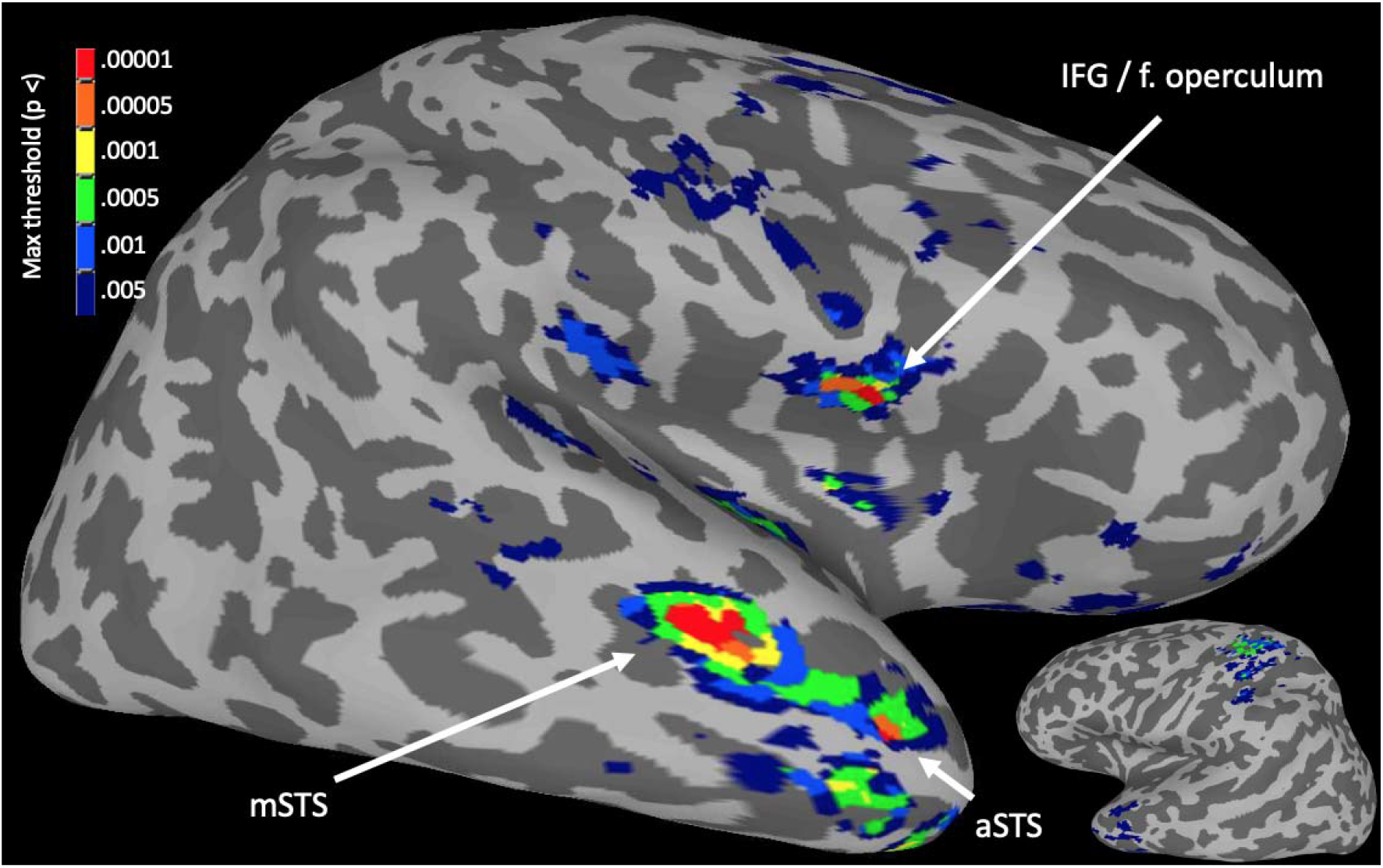
Right IFG, right mSTS, and right aSTS showed greatest Autism > Typical whole-brain connectedness across statistical thresholds. Inflated surfaces of the right hemisphere (large) and left hemisphere (small, inset). Colors indicate most stringent threshold survived.

For each of these three target ROIs, we assessed the correlations between wholebrain functional connectivity and Classifier Score by again fitting linear models with whole-brain connectedness (“Connectivity”) predicted by Classifier Score, Group (Autism or Typical) and their interaction. The interaction effect, critical for assessing compensation, was tested for each of the ROIs at a Bonferroni-corrected threshold of *P* < .05 / 3 tests = .0167.

### Data availability

The LIWC output and ROI-level neuroimaging data used in the analyses will be available on publication.

## Results

First, we assessed the accuracy and validity of the behavioral measure by evaluating how well the classifier could distinguish Autism and Typical participants when trained on participants’ speech. As a control, we also assessed whether the classifier could distinguish Autism and Typical participants if trained on the experimenter’s speech. External validity was assessed by comparing how the classifier scored autism participants during the interactions to how a clinician had scored the same participants using the ADOS. Finally, the relationship with functional connectivity was assessed. If elevated functional connectivity is compensatory, it should correlate with typical behavior for the Autism participants, but not for Typical participants (who should have no need to compensate).

### Classifier accuracy and validity

The overall accuracy of the classifier trained on participant speech was 74% [*P* = .002 by permutation test; Fig. 2A]. Autistic participants with more Typical classifier scores also displayed fewer autistic signs during their diagnostic observation performed a clinician [correlation with ADOS-2 Social Communication scores, Spearman *r* = –.51, *P* = .03; Figure 2B]. It is of note that the two autism participants who were misclassified as typically developed by the classifier had the minimum ADOS-2 Social Communication scores required for diagnosis (i.e., a score of 7).

**Figure 2.**
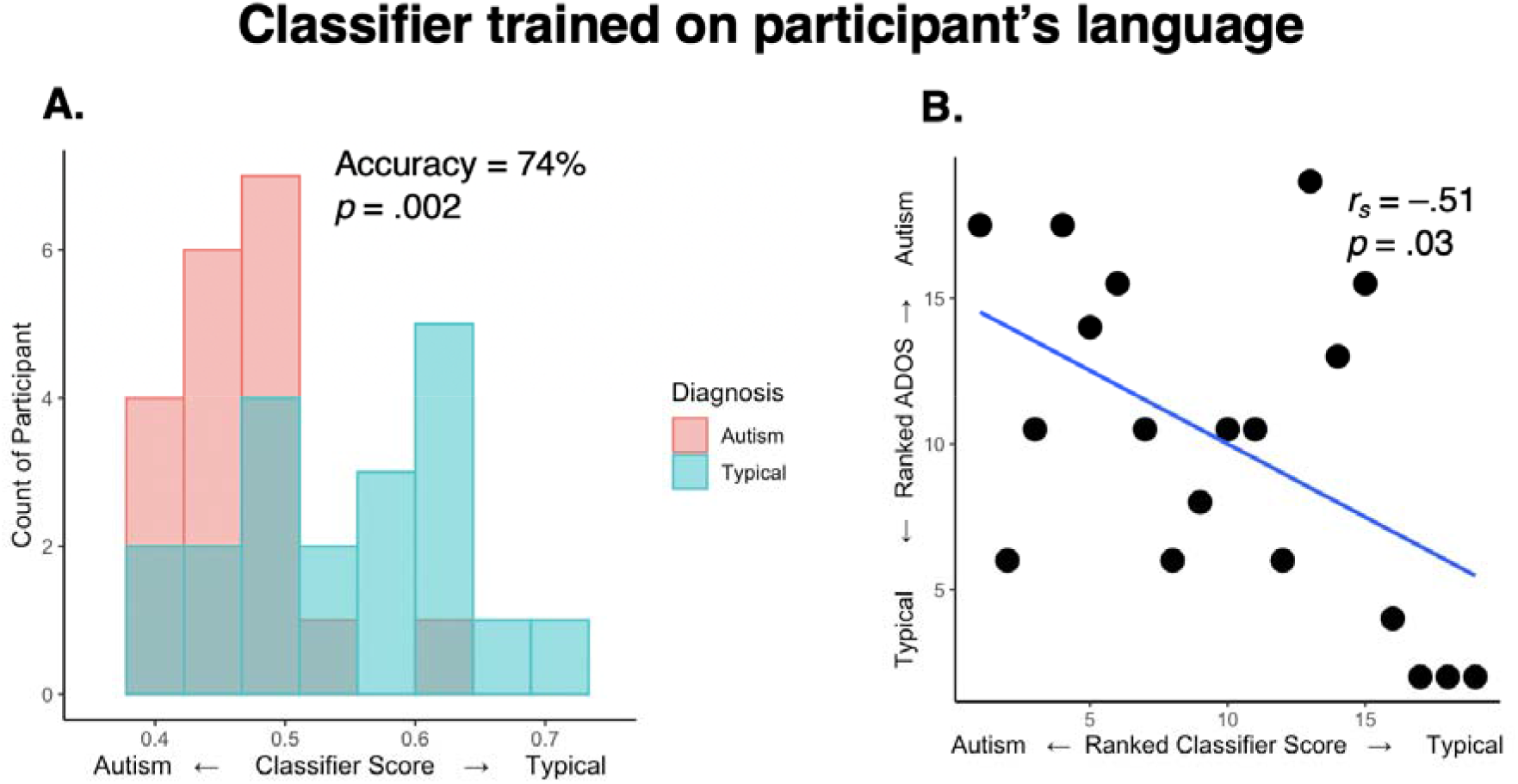
Classifier scores distinguished Autism and Typical participants and correlated with ADOS-2 Social + Communication scores. **(A)** Histogram of mean classifier scores by diagnostic group and **(B)** clinician ADOS-2 Social Communication subscores of autism symptom severity plotted by ranked classifier score.

As a control, the same classification analyses were performed on the experimenter’s speech. Classifier accuracy dropped to 59%, which was statistically indistinguishable from chance [*p* = .13 by permutation test; Figure S1.] The scores derived from the experimenter’s speech furthermore did not correlate with ADOS-2 scores [Spearman *r* = –.20, *P* = .42 Fig. S1]. ROC curve analyses also indicated good classification when using participant speech, but poor classification when using experimenter speech (Figure S2).

### Typical language behavior is associated with elevated functional connectivity

Having established the validity of the linguistic measure, we examined its relationship with functional connectivity during the sessions. Three regions of interest were examined: mSTS, aSTS and RIFG. For each of these, the interaction of Group with Classifier Score was tested. Only the test for RIFG survived Bonferroni correction. (Right mSTS survived an uncorrected threshold of P < .05; for transparency we report this analysis in the Supplement (Figure S4), and the pattern was qualitatively similar to that found in the RIFG.) For RIFG, the slope of the correlation between functional connectivity and classifier score differed between the two groups (interaction *t*(35) = –2.7, *p* = .01; Figure 3): in the Autism group, more typical classifier scores were associated with higher functional connectivity (*r*(17) = .56, *p* = .01), but this pattern failed to be detected in the Typical group (*r*(17) = –.18, *p* = .44; Figure 3).

**Figure 3.**
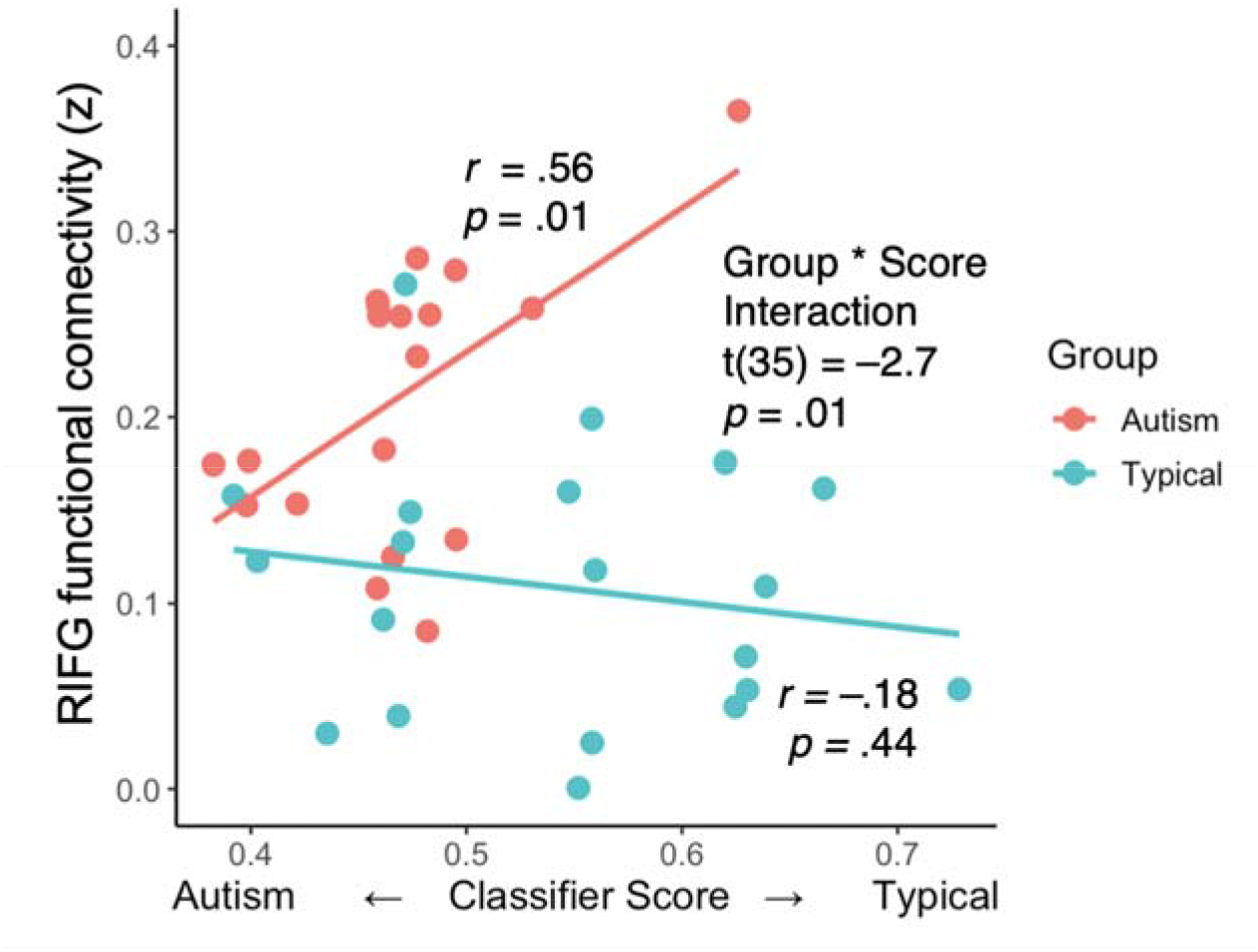
Typical language production was correlated with higher RIFG connectivity in autism participants. Scatter plot of whole-brain connectivity of RIFG plotted against Classifier Score for the autism (red) and typical (teal) groups.

Our neural measure, average “connectedness” with the whole brain, gives a broad indication of the level of functional involvement of a given region (in this case RIFG). However, the specific pattern of connectivity with the RIFG is unclear without followup analyses. We therefore ran an exploratory seed-based functional connectivity analysis using the RIFG to determine which other regions were involved in overconnectivity. A group comparison (Autism > Control, P < .001, corrected) was performed after fitting a linear mixed effects model to the data with *3dLME* ([Connectedness ~ Group + Age + Motion + (1 | Participant)]). The results indicated that more than twice as many left-hemisphere voxels were over-connected with RIFG (5317 voxels) compared to right-hemisphere voxels (2452 voxels). The most prominent results were in left hemisphere areas associated with language and social interaction, such as left inferior frontal gyrus, left superior temporal gyrus and temporal pole, and left temporo-parietal junction (Figure 4). The compensatory effect of right IFG connectivity appears to be driven strongly by its coordination with contralateral left hemisphere language areas.

**Figure 4.**
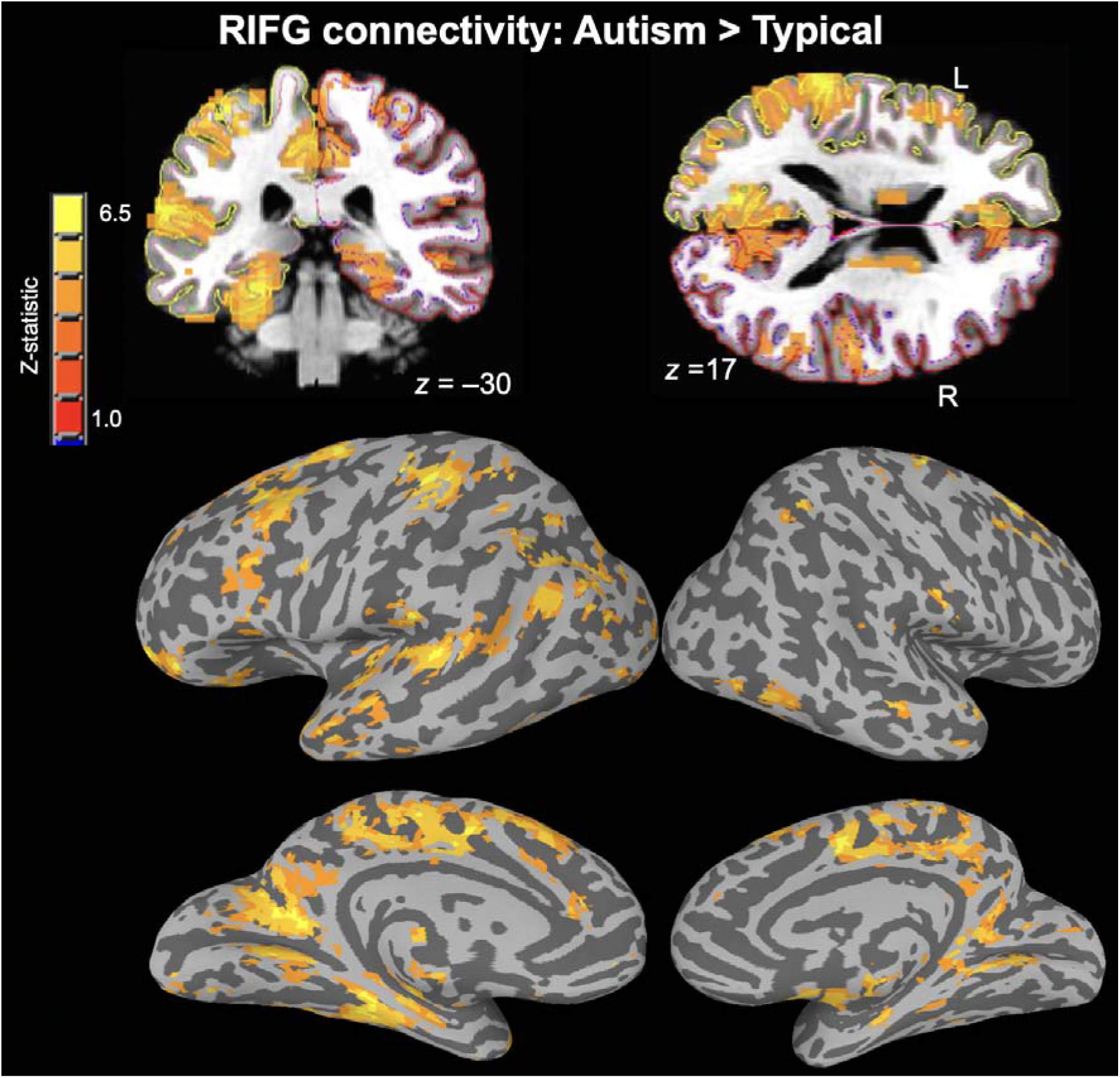
Compensation takes place through cross-hemispheric connectivity between left hemisphere social communication areas and right IFG. Coronal and axial sections, and inflated surfaces showing clusters with significant Autism > Typical functional connectivity with RIFG during conversation (P < .001, corrected). More than twice as many significant voxels were in the left hemisphere, and the strongest results were in regions involved with social communication and language.

## Discussion

In a previous study we detected widespread functional connectivity increases in autism during conversation. Here we found that autism participants with higher functional connectivity, particularly involving the RIFG, produced more typical language during conversations. This result helps to clarify the relationship between functional connectivity and performance of a naturalistic social task in autism, namely that functional connectivity increases are in some cases compensatory, helping rather than hindering social communication.

Why did RIFG emerge as a particularly important region for compensation? Conversation of course relies heavily on speech and language production, and our behavioral measure was derived entirely from spoken language. Although language is typically left-lateralized, there is evidence that the contralateral right hemisphere homologs of left-hemisphere language areas are recruited when task demands are high, both in typically developed individuals and patient groups (so called ‘spillover’ processing, Prat et al., 2011). Relating to autism and RIFG specifically, activity in RIFG (the homolog of Broca’s area) has been shown to increase when autism participants integrate language with social information such as age and gender (Tesink et al., 2009). Under this ‘spill-over’ account, the reason RIFG is heavily involved is that conversation is difficult for autism participants, and contralateral LIFG cannot meet task demands on its own. Our results are also consistent with a recent paper by Persichetti and colleagues, who found that the relative lack of cross-hemisphere interactions from right-hemisphere language homolog regions in autism was associated with poorer verbal ability (Persichetti et al., 2022). A second and not-necessarily mutually exclusive possibility is that RIFG is playing a role in ‘executive function’, a process that has been proposed to assist compensation in developmental disorders (Johnson et al., 2015).

The results do not fully address the question of which patterns of connectivity reflect underlying deficits and which may have developed to compensate for deficits. Further work should clarify this by quantifying other aspects of behavior and relating this to ongoing neural connectivity. The present study only used counts of spoken words in various categories as raw data. Other studies could examine, for example, acoustic aspects of vocal production such as prosody, eye movements, or hand gestures. Still other studies might address more subtle communication abilities such as how theory of mind problems are solved.

Further work should investigate a more diverse set of participants. Participants in this study had normal to relatively high IQ. It remains to be seen whether individuals with lower IQs also compensate, and if they do, whether they use a similar or different neural strategy. It has been suggested that females with autism may compensate more successfully than males (Dworzynski et al., 2012). Future studies could concentrate on other populations such as female participants, or perhaps so-called ‘unaffected’ siblings of individuals with autism, who may exhibit only mild autistic traits and receive no diagnosis due to a covert neural compensation strategy (Johnson et al., 2015).

A minor result from Jasmin et al., 2019, was that RIFG connectivity with R parahippocampal gyrus was positively correlated with SRS, suggesting that functional connectivity increases correlate with more severe core autism deficits. That result is not necessarily inconsistent with the results reported here. As mentioned in the introduction, tools like the SRS measure persistent core autism deficits, while “one-off” social situations such as an ADOS-2 interview, or the one-to-one conversation task in this study, may present optimal conditions for compensation (Livingston & Happé, 2017). Indeed, SRS and ADOS-2 scores exhibit little or no correlation (Morrier et al., 2017). It is therefore possible that measures of autism “trait” vs. current behavioral “state” may correlate with connectivity differently.

The classifier-based behavioral method described here has wide applicability for studying compensation in developmental disorders. A main benefit is that it is not necessary to have an *a priori* definition of typical behavior. Behavior of any two groups of participants who are matched for relevant variables but differ categorically in diagnostic category could be submitted to a classification algorithm, and the resulting probability score could be used as an index of behavioral ‘performance’ and linked with neural measures.

## Acknowledgements

We thank our participants and everyone who contributed to this project.

## Supplementary material

Supplementary material is available online.

