## Supplement for "Compensation and conversation in autism: atypical connectivity supports typical behavior"

### Supplementary Materials for Jasmin, Martin, & Gotts, 2023.

**Table S1 Region of Interests**

| Anatomy | X | Y | Z | *N* voxels | Peak *Z* |
| --- | --- | --- | --- | --- | --- |
| R IFG | 44 | 5 | 8 | 10 | 5.1 |
| R mSTS | 47 | -16 | -10 | 23 | 5.0 |
| R aSTS | 53 | 8 | -19 | 10 | 4.7 |

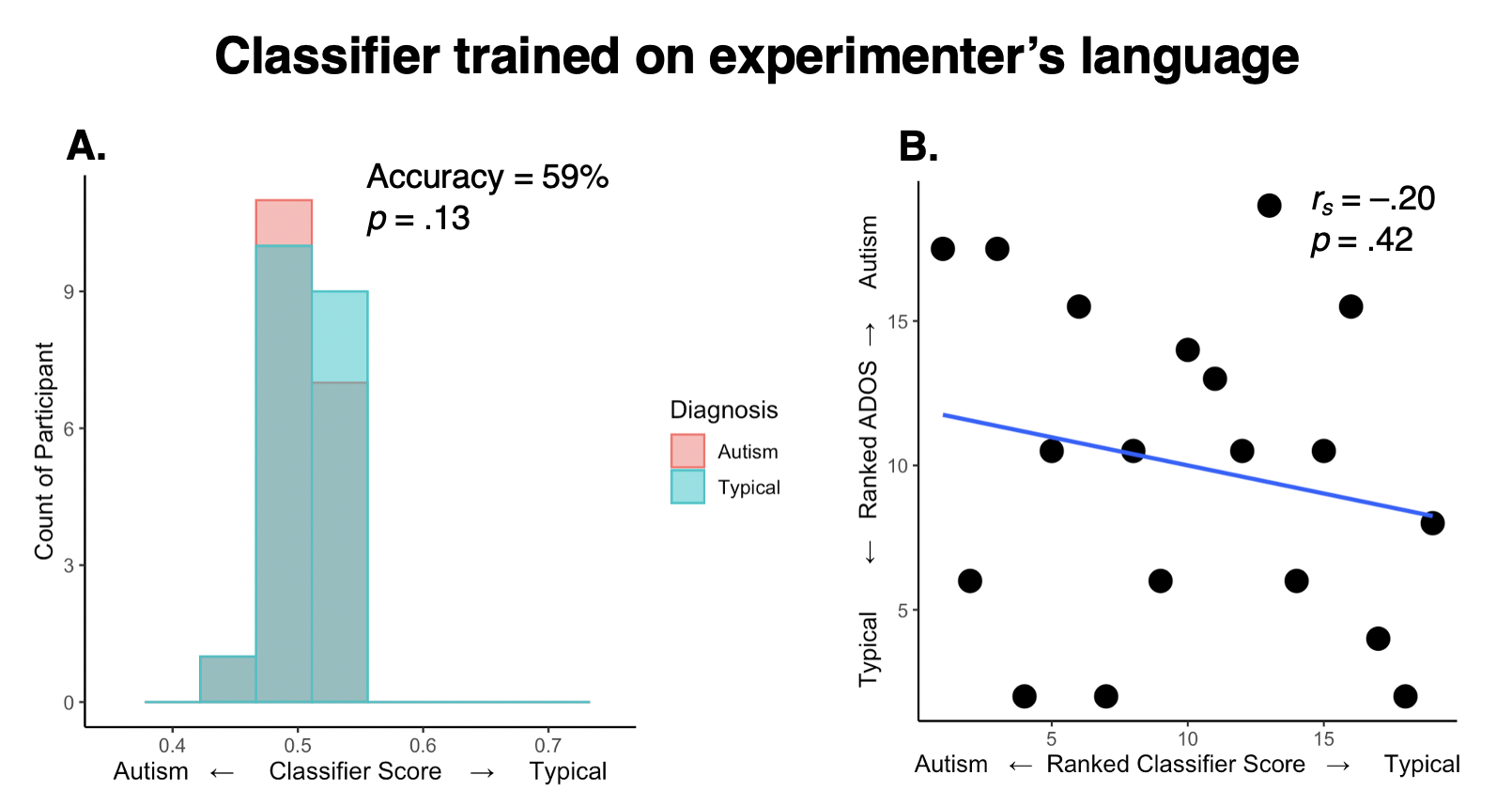

**Figure S1 The classifier trained on experimenter speech neither distinguished Autism and Typical participants nor correlated with ADOS-2 Social + Communication scores.** For the classifier trained on experimenter’s speech, **(A)** histogram of mean classifier scores by diagnostic group and **(B)** clinician ADOS-2 Social Communication subscores of autism symptom severity plotted by ranked classifier score.

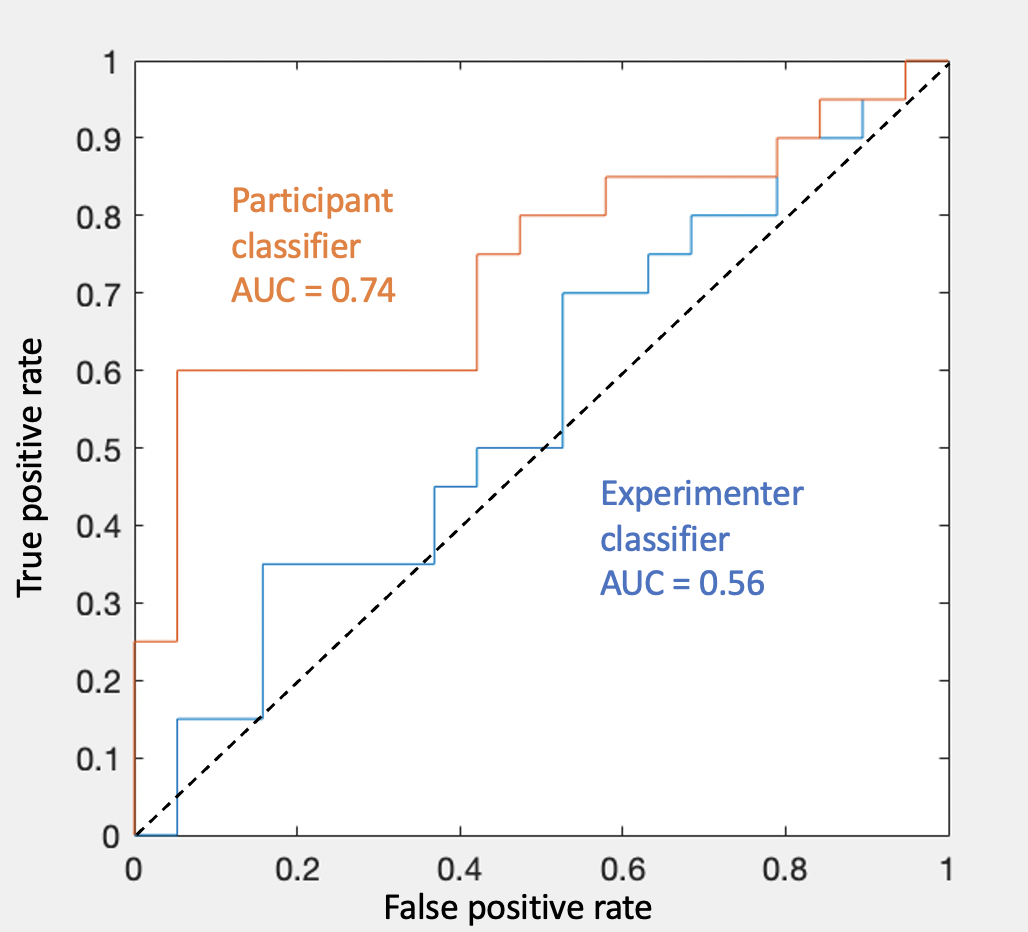

**Figure S2 Receiver Operator Characteristic curves for the participant-trained classifier and the experimenter-trained control classifier** AUC = Area Under the Curve. Values give an indication of proportion of times a randomly selected Classifier Score value from the Typical group would have a higher value than a randomly selected value from the Autism group. Dashed line indicates chance classifier performance. In terms of effect size an AUC value of 0.74 is equivalent to Cohen’s *d*  = 0.9 while an AUC of .56 is equivalent to *d* = 0.2 (Rice and Harris 2005).

**Analysis of Classification Features**

An analysis of the text was undertaken to examine which features of the text were associated with classification scores. First, across all 115 conversations the classifier scores were correlated with the *z*-scored linguistic (LIWC) features. Statistical significance was assessed at FDR(*q*) < .05, and for transparency, also at uncorrected *p* < .05. The relative magnitude of the feature weights of the SVM classification (*β*^2^) was also examined to confirm that features with strong correlations with classification probability were also weighted heavily in the model.

After correlating the 81 LIWC features with the classifier scores, 15 features showed a significant (*p* < .05) association with the scores (Figure S2A). Only one of these categories, “Linguistic Conjunctions” (e.g. “and”, “so”, “but”), survived False Discovery Rate (FDR) correction for multiple comparisons, and was associated with higher probability of classification as typical development (*r* = .38, *p* < .001). This category was also associated with the largest feature weight (*β*^2^ = 2.73). To ensure the pattern was not simply driven by an overall group difference, the count of linguistic conjunctions (*z*-scores) and classifier scores were averaged by participant, and correlations were examined separately for each group. Use of conjunctions was associated, separately, with Typical classification in both the autism group (*r* = .50, *p* = .03) and typical group (*r* = .45, *p* = .04; Figure S1).

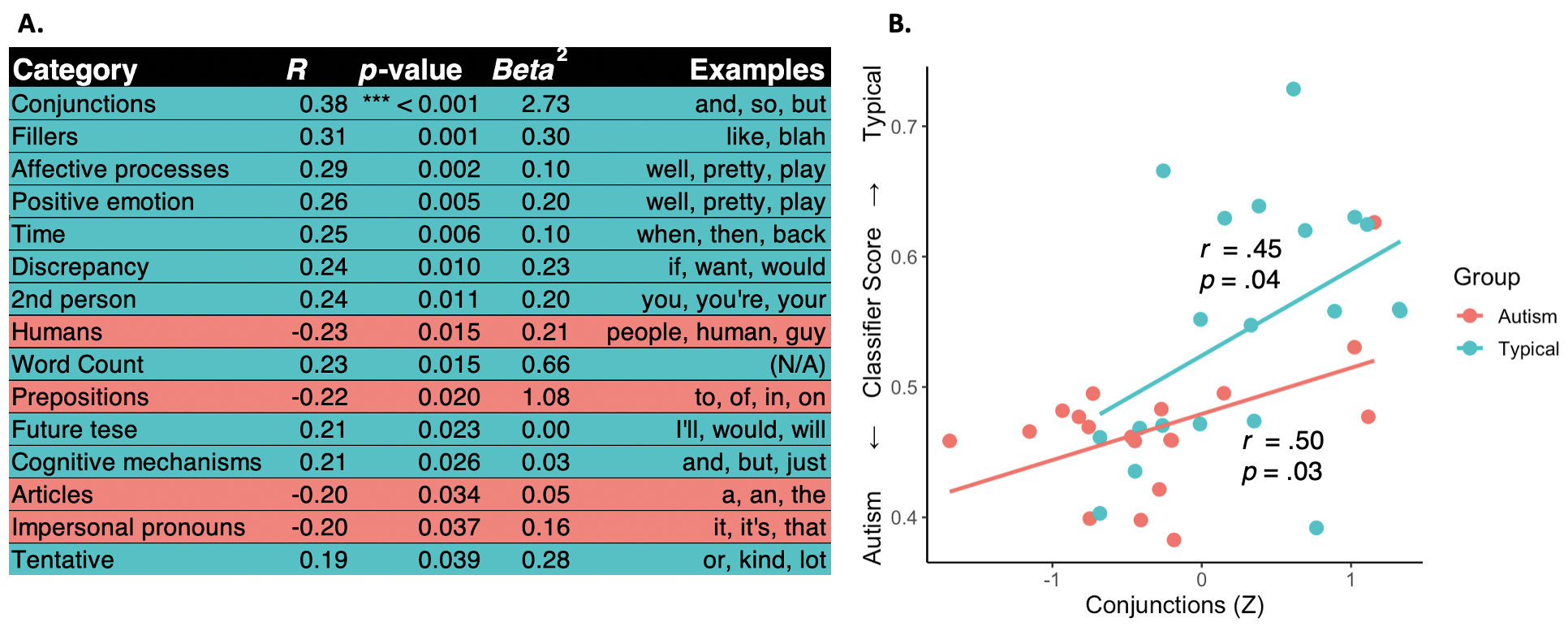

**Figure S2 Linguistic features associated with typical vs. autism classification. A)** LIWC categories sorted by p-value. Blue highlights indicated positive correlation with Typical classification; Red highlights indicate negative correlations. Beta^2 is provided as a measure of feature weight. Only the Conjunctions category survived correction for multiple comparisons. **B)** Plot of classifier score by count of conjunctions (Z-scored), by participant. Greater use of Conjunctions was associated with greater probability of Typical classification, for both the autism and typical groups.

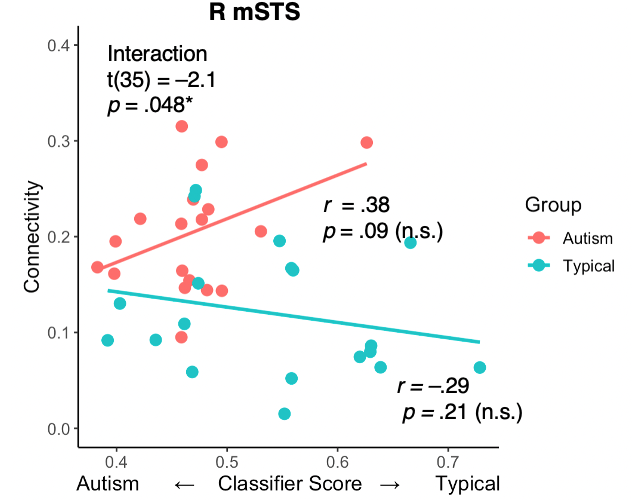

**Figure S4 The relationship between Classifier Score and R mSTS-to-whole-brain connectivity by Group (p<.05).** The interaction effect did not survive the Bonferroni-corrected threshold but is reported here for transparency. The signs of the correlations are overall similar to those of the R IFG analysis, although the relationship between Score and Connectivity in the Autism group was not significant even at P < .05.
